# Neq2X7: a multi-purpose and open-source fusion DNA polymerase for advanced DNA engineering and diagnostics PCR

**DOI:** 10.1101/2022.03.14.484273

**Authors:** Cristina Hernández-Rollán, Anja K. Ehrmann, Morten H. H. Nørholm

**Affiliations:** The Novo Nordisk Foundation Center for Biosustainability, Technical University of Denmark, Kemitorvet Building 220, 2800 Kongens Lyngby, Denmark; Mycropt ApS, 2800 Kongens Lyngby, Denmark

**Keywords:** *Nanoarchaeum equitans* Neq DNA Polymerase, Polymerase Chain Reaction, modified nucleotides, uracil-excision cloning, USER cloning, dUTP

## Abstract

Thermostable DNA polymerases, such as Taq isolated from the thermophilic bacterium *Thermus aquaticus*, enable one-pot exponential DNA amplification known as polymerase chain reaction (PCR). However, other properties than thermostability - such as fidelity, processivity, and compatibility with modified nucleotides - are important in contemporary molecular biology applications. Here, we describe the engineering and characterization of a fusion between a DNA polymerase identified in the marine archaea *Nanoarchaeum equitans* and a DNA binding domain from the thermophile *Sulfolobus solfataricus*. The fusion creates a highly active enzyme, Neq2X7, capable of amplifying long and GC-rich DNA and that is unaffected by replacing dTTP with dUTP in PCR. This makes it an attractive DNA polymerase for use e.g., with uracil excision (USER) DNA assembly and for contamination-free diagnostics. Furthermore, Neq2X7 is easy to produce, and the expression plasmid is freely available.

**Graphical abstract:** 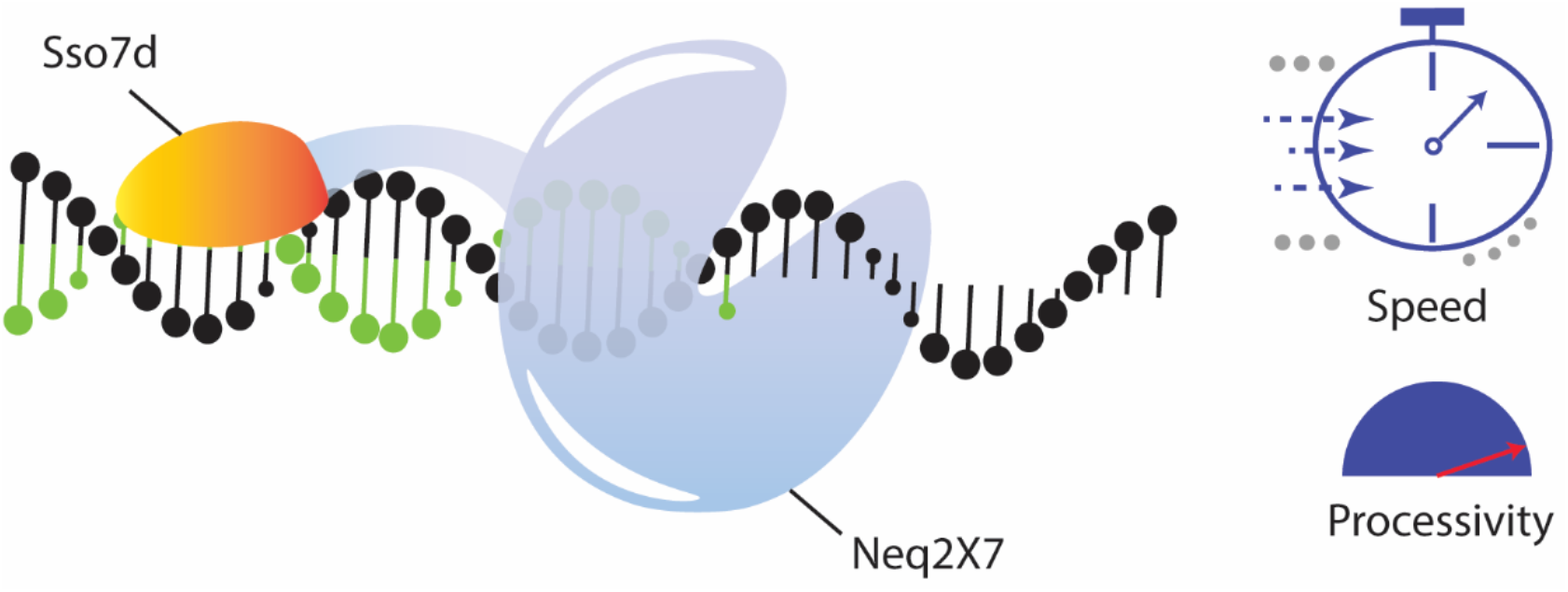

## Introduction

Exponential DNA amplification by the polymerase chain reaction (PCR) is one of the most important inventions in molecular biology, and the technique was paradigm-shifting in DNA diagnostics, and forensic science^1^. DNA polymerases are the central enzymes in PCR and are at the frontier of other biotechnological applications^2,3^. Many contemporary synthetic biology applications rely entirely on PCR, and DNA polymerase properties like high-fidelity and processivity are imperative. Applications that require exceptional DNA polymerases include advanced DNA assembly methods^4–10^, medical diagnostics^11^, or technologies that require the incorporation of non-conventional nucleotides such as xeno-nucleic acids (XNA)^12^. Additionally, next-generation DNA sequencing approaches are continuously refined and rely on DNA polymerases that tolerate specifically labeled substrates^13^. Finally, routine PCR workflows that allow for amplification of DNA in short time frames - as low as 3 minutes^14^ - are crucial for rapid detection of e.g., pathogens, and are particularly relevant during pandemic events, such as the current Covid-19 crisis.

Most PCR methods rely on the action of a thermostable DNA polymerase with high processivity and low error rate^15^. Depending on the specific applications, other desired features include high DNA yield, and amplification of long (long-range PCR) or complex templates (e.g., DNA with high GC content or secondary structure). In specific cases it is a requirement that the DNA polymerase tolerates a wide range of nucleotides such as deoxyuridine triphosphate (dUTP). The popular DNA assembly method USER cloning^6,16,17^ utilizes uracil bases in the PCR primers to create short compatible single-stranded overhangs. Incorporation of dUTP into the DNA template also plays a role in another PCR application since it can be used to limit carry-over of amplification products in sensitive environments such as forensic laboratories^18^. When PCRs are performed using dUTP instead of dTTP, reactions can be treated with uracil-N-glycosylase (UNG) before amplification. This way all uracil-containing DNA fragments are degraded and only the test template is left intact^19^. Finally, since some uracil is found in environmental DNA, uracil-accepting polymerases should amplify such DNA more robustly^20^.

Archaeal DNA polymerases, mainly belonging to family B, are ideal candidates for PCR, as they share most of the crucial characteristics for advanced applications: they are often thermostable and show high-fidelity thanks to their proofreading (3’-5’-exonuclease) activity^21^. Furthermore, they can amplify long stretches of DNA and, in contrast to Taq polymerase, they do not add an extra dA at the end of the amplification. These characteristics make them highly attractive for PCR and cloning techniques. Still, a wildtype DNA polymerase rarely possesses all the ideal characteristics needed and protein engineering is often carried out to improve the performance of natural DNA polymerases^13^.

In 2006, the expression and characterization of a new archaeal DNA polymerase from the hyperthermophile *Nanoarchaeum equitans* (Neq) was reported^22^. Later, the use of Neq for PCR was described and it was discovered that, in contrast to other archaeal family B DNA polymerases, Neq does not stall when encountering uracil, as it lacks the specific binding pocket^23,24^. Additionally, Neq has high processivity but low fidelity. However, the fidelity was improved through the incorporation of two mutations (A523R/N540R) which brought the error rate of the Neq polymerase *on par* with the highly popular Pfu polymerase^25^. Nevertheless, despite its apparent attractive properties as a high-performance polymerase for *in vitro* applications, Neq does not yet seem to be widely used as judged by a simple internet keyword search (Supplementary Figure 1).

There are different ways to engineer a given DNA polymerase for enhanced processivity. One of the simplest approaches reported to date is the attachment of an unspecific DNA binding domain^26^. The DNA binding domain of *Sulfolobus solfataricus* (known as Sso7d domain) enables higher processivity through its dsDNA binding ability without significantly changing the other catalytic properties of the polymerase^3,7^.

Here we engineer and characterize a new variant of the Neq polymerase, Neq2X7, that combines the two fidelity-increasing mutations (2X) previously reported with the addition of Sso7d^27^. The engineered polymerase shows improved processivity over Neq(A523R/N540R) (Neq2X) and PfuX7 and perform well in amplification of DNA up to 12000 bp, in high GC content DNA amplification, and in incorporation of dUTP. Neq2X7 is an efficient and versatile polymerase suitable for routine molecular biology as well specialty applications, like USER cloning, or prevention of PCR carry-over. Finally, Neq2X7 is easily produced with a routine protein production workflow and the construct is freely shared with the community and accessible via the Addgene repository.

## Results and Discussion

### Expression plasmid construction, DNA polymerase expression, and purification

The double mutant Neq2X gene was ordered as a synthetic gene and cloned into two different pET plasmids for production in the standard *Escherichia coli* pET system with or without the Sso7d-encoding domain as previously described for Pfu^7^. The resulting DNA polymerases carry 6xHis purification tags at their N-termini, and Neq2X7 harbors the Sso7d DNA fusion domain at the C-terminus (Figure 1A). Using a typical and simple protein production protocol (see methods and Supplementary Figure S2), we estimate that we produce enough Neq2X7 DNA polymerases for about 50,000 PCR reactions starting out with a standard 100 mL of bacterial batch culture. The constructs are summarized in a table in Figure 1B along with Addgene repository accession numbers^28^.

**Figure 1.**
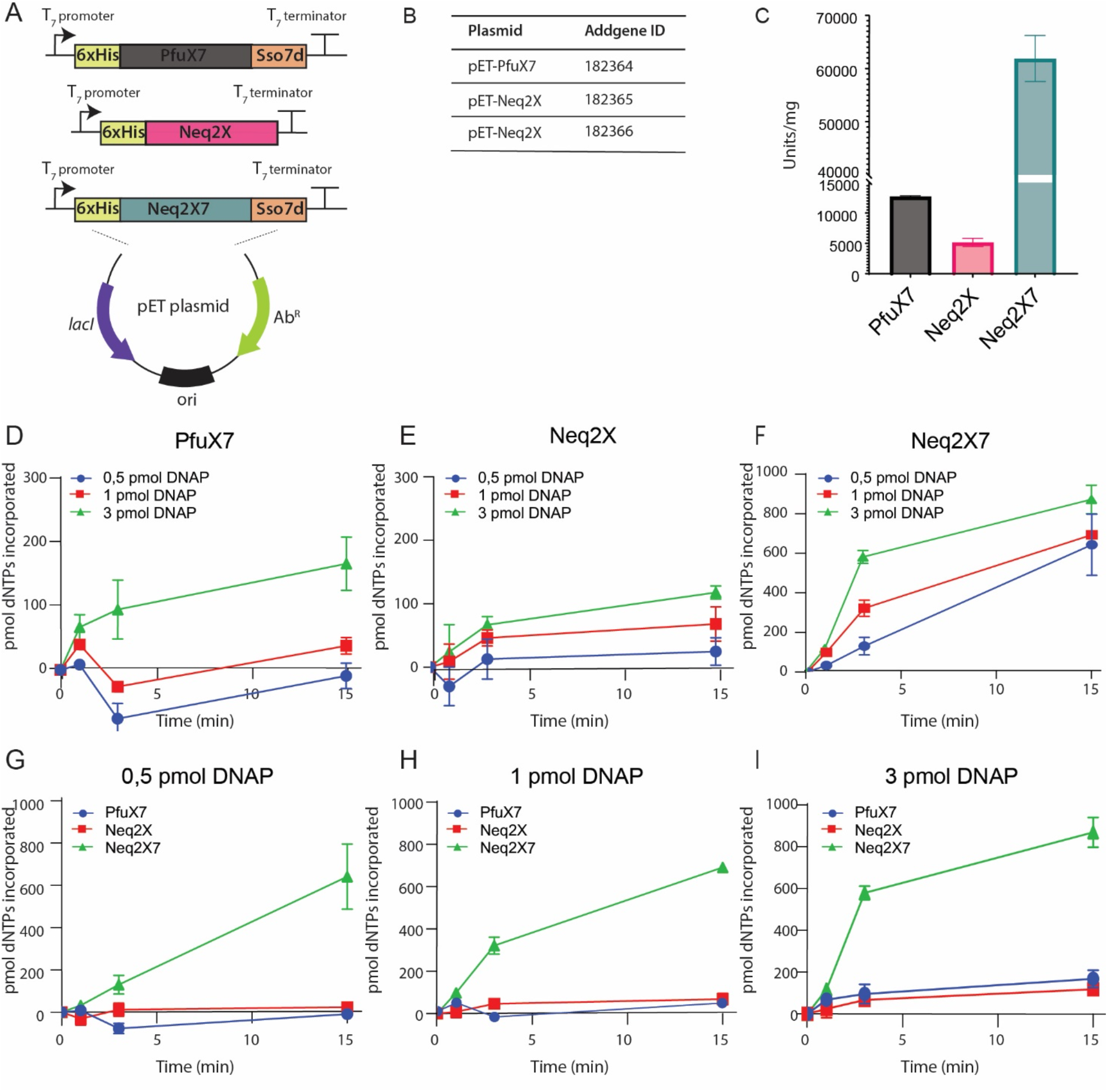
Illustration of the plasmids used in this study, and comparison the PCR extension rates of the DNA polymerases PfuX7, Neq2X and Neq2X7 measured by binding of the fluorescent Pico Green dye to double stranded DNA. A) Schematic overview of the PfuX7, Neq2X, and Neq2X7 expression plasmids. B) Table with plasmids used in this study including their Addgene IDs. C) DNA polymerase activity units calculated based on incorporation of dNTPs in 10 minutes per mg of purified enzyme. D) Fluorescence-based activity assay for PfuX7, E) Neq2X, and F) Neq2X7 using 0.5, 1, and 3 pmol of enzyme over time. G) Fluorescence-based activity assay comparing dNTP incorporation rates between PfuX7, Neq2X, and Neq2X7 for 0.5 pmol, H) 1 pmol, and I) 3 pmol of each enzyme and over time. All measurements were performed in triplicates, and the graph displays the median with the standard deviation indicated. DNAP (DNA Polymerase). PicoGreen measurements were done using a Synergy H1 plate reader (BioTek Instruments, Inc).

### Benchmarking the activity of Neq2X7 with similar DNA polymerases

The performance of Neq2X7 was compared with the mutant Neq polymerase without the Sso7d DNA binding domain (Neq2X) and the Pfu-derived Sso7D fusion DNA polymerase PfuX7^7^ using a standard fluorescence-based DNA polymerase assay. This analysis showed a high activity of Neq2X7, compared to the other two polymerases, as reflected in the measured units per mg of the enzyme (Figure 1C), which was extrapolated from the initial time points where the rate of incorporation of dNTPs progresses in a linear fashion (Figure 1D-I) and was normalized by the amount of enzyme. Looking closer at the data, using three different protein concentrations (0.5, 1, and 3 pmol), and time points after 1, 3, and 15 min, it can be observed that Neq2X7 incorporates more dNTPs on shorter timescales, even with less enzyme present. While polymerase activity is detected with as low as 0.5 pmol for Neq2X7 (Figure 1F), PfuX7 and Neq2X only show detectable incorporation of nucleotides at 3 pmol (Figure 1D and 1E). Units are typical measures of enzyme activity and are defined here as the amount of polymerase that incorporates ten pmol of dNTPs using a primer together with single-stranded viral DNA from M13mp18 as a template at 72°C in 10 minutes. The activity of the Neq2X7 increases about eight-fold with the addition of the Sso7d DNA binding domain.

### Benchmarking the performance of Neq2X7 in various PCR applications

One of the classical challenges of PCR is amplification of very long DNA stretches since this increases the risk of premature replication termination and this is why highly processive DNA polymerases are desirable^21^. We compared Neq2X7, Neq2X and PfuX7 PCR performance using three different amplicon sizes: 3, 6 and 12 kb. With an extension time of 1 min/kb, all three polymerases managed to amplify DNA to a detectable level (Figure 2A). However, when the extension time was shortened to 15 sec/kb, only Neq2X7 was able to produce the desired DNA fragments (Figure 2B). This indicates that the processivity of Neq2X7 is high compared to Neq2X and PfuX7.

**Figure 2.**
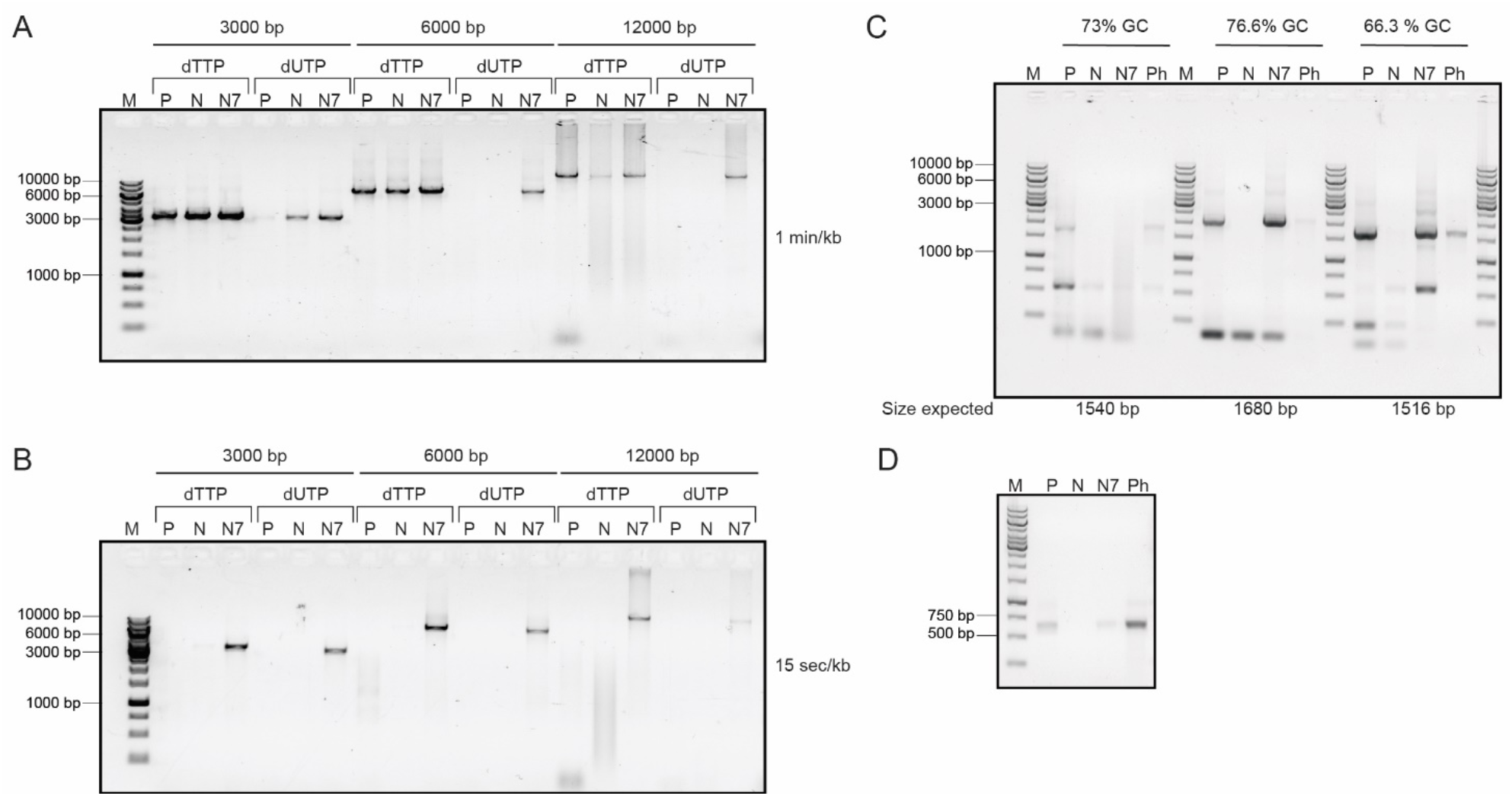
PCR comparison of the DNA polymerases performance for different templates, amplification times, and conditions using 1pmol of each DNA polymerase enzyme run on a 1% agarose gel. A) Three different templates ranging from 3000, 6000, and 12000 bp were amplified using 1 minute of amplification per kilobase. For each template two conditions were evaluated, (i) the amplification of DNA with a mixture of dNTPs containing **dTTP** (dATP, dTTP, dCTP, and dGTP), and (ii) **dUTP** replacing dTTP (dATP, dUTP, dCTP, and dGTP). B) The same templates as in panel A were amplified using 15 seconds of amplification time per kilobase. C) High GC content templates amplification from three different genomic DNA regions of Streptomyces were amplified by PCR. From left to right, 73% GC of 1540 bp expected size, 76.6% GC of 1680bp expected size, and 66.3% GC of 1516 bp expected size. D) Fast-PCR amplification of a 630 bp template. P (PfuX7), N (Neq2X), N7 (Neq2X7), and Ph (Phusion) DNA polymerases. M) DNA marker in base pairs. For PCR conditions, primers, and templates, see the methods section.

A normal characteristic of archaeal DNA polymerases is the presence of a conserved domain called the uracil-binding motif^21^. This domain recognizes uracil deaminated bases, and upon binding, DNA amplification stalls. It is believed that this domain evolved to protect the archaeal organisms from uracil incorporation that results in the deamination of cytosine, which can be mutagenic. However, the Neq polymerase lacks this uracil-binding pocket and therefore has the natural ability to tolerate uracil in the template and PCR amplification with dUTP nucleotides. In contrast the “X” in PfuX7 denotes an engineered mutation that compromises the uracil-binding activity of this polymerase. We compared the performance of Neq2X7, Neq2X and PfuX7 in PCR amplification with dUTP entirely replacing dTTP in the dNTP mix. For the shorter amplicon of 3000 bp, all tested polymerases were able to generate a detectable PCR product with an extension time of 1 min/kb (Figure 2A). For longer amplicons, and at the reduced extension time of 15 sec/kb, only Neq2X7 was able to amplify the target DNA successfully with dUTP (Figure 2B). This indicates that Neq2X7 is a superior polymerase for applications requiring dUTP.

Often, a limitation in the use of DNA polymerases in PCR is the amplification of DNA with high GC content. Many biotechnologically interesting organisms, such as *Streptomyces*, and mycobacteria ^29,30^ - and sometimes specific genomic regions ^31^ - have high GC content and therefore are challenging to work with. We evaluated the performance of Neq2X7 on high GC content templates using three different templates from genomic DNA isolated from *Streptomyces* ranging from 66 – 76 % GC content (see Supplementary Table 4). While Neq2X was unable to amplify any of the targets, the correct product was detected for Neq2X7 in two out of three cases (Figure 2C). PfuX7 and the commercially available Phusion DNA polymerase (Invitrogen) were able to amplify the correct product for all three reactions. Thus, the Pfu polymerases may be better choices for high GC content DNA. It is perhaps not surprising that Neq has difficulty handling high-GC content templates since the GC content of the genome of *N. equitans* is only 31.6 %. Still, the addition of the Ssod7 domain considerably improves the ability of Neq to amplify high-GC DNA.

Fast DNA amplification is important for diagnostics and allow for amplification of DNA targets in the range of minutes. Inspired by the promising results on the high processivity of Neq2X7, we wanted to test whether this improved polymerase can perform “Fast-PCR”, and how its performance would compare to other highly processive polymerases (e.g., PfuX7 and Phusion). A 653 bp DNA fragment was selected for amplification using a fast-PCR program with a total runtime of 24 minutes (Supplementary Table 5). Both PfuX7 and Neq2X7 show a detectable signal on the agarose gel after the PCR, even though they are outperformed by the commercially available Phusion polymerase (Thermo Scientific). Further optimization of the protocol and the buffer conditions, as well as utilization of specific Fast-PCR equipment, might enable Neq2X7 to compete head-to-head with the fastest PCR protocols currently available.

A common strategy to increase the efficiency of PCR reactions is to mix Taq polymerase with archaeal family B DNA polymerases. This strategy was previously employed to enhance the performance of the Neq polymerase^23^. We tested the combination of Neq2X7 and DreamTaq (Thermo Scientific) in two different concentrations to amplify two different targets (Supplementary Figure 3), but did not observe considerable improvements in DNA amplification. Instead, there is an increase in unspecific amplification products if Neq2X7 is either mixed with DreamTaq, or if the concentration of Neq2X7 is increased. DreamTaq alone was not able to amplify the targets in the applied reaction conditions. These results highlight that adding more polymerase does not always result in better results, and that mixtures of different polymerases have to be carefully tuned to achieve the desired outcomes. Other polymerase combinations or further optimization of the reaction conditions could be interesting to test with Neq2X7.

## Conclusions

The addition of the Sso7d binding domain to the double mutated Neq DNA polymerase yields a highly active DNA polymerase, Neq2X7, that in the last couple of years has proven superior to most other available DNA polymerases in our laboratory. Here, we demonstrate this for long template and short extension time PCRs and with dUTP replacing dTTP entirely in the dNTP mix commonly used in PCR. In a synthetic biology laboratory like ours, the latter is of particular interest in combination with the DNA assembly method known as USER cloning, but the enzyme could also find good use in forensic laboratories utilizing dUTP incorporation for prevention of template carry-over.

The observation that Neq2X7 outperforms and enzyme like Pfu in some of the PCR applications tested here is an indication that the natural capabilities to accept uracil containing DNA is an attractive feature that should be further investigated in other thermostable organisms. The Neq2X7 expression construct is available through the DNA repository Addgene and the enzyme is easy and cheap to produce and purify using standard protocols. We hope that the sharing will facilitate its use in creating useful synthetic biology designs and biotech applications or in relevant teaching environments with limited resources.

## Methods

### Vector constructions

We used the (in-house) pET-PfuX7 vector expressing PfuX7 DNA polymerase as described in^7^. A pET-vector expressing Neq2X DNA polymerase^25^ with the Sso7d DNA binding domain fused to the polymerase C-terminus (See Figure 1A) was synthesized by GeneScript and we named it Neq2X7. The Neq2X DNA polymerase vector was created by removing the Sso7d DNA binding domain by USER cloning, with primers: 4369 and 4370 as described in ^6^ and cloned into *E. coli* NEB5α (New England Biolabs, Ipswich, MA, USA) competent cells.

### Protein production

All three vectors were transformed into chemically competent *E. coli* Rosetta2DE3pLysS (Novagen, Merck, KGaA, Darmstadt, Germany). A single colony derived from the transformation was used to set an overnight culture in a 3mL 2xYT medium at 37 °C with 250 rpm of shacking. Cultures were supplemented with ampicillin and chloramphenicol for PfuX7 plasmid and kanamycin and chloramphenicol for Neq2X and Neq2X7 plasmids. The next day, the overnight culture was diluted 1:100 into 100 mL 2xYT media with the same antibiotics and growth conditions. When the culture reached an OD600 of 0.3, the expression was induced with 0.5 mM of IPTG. After four hours, the cultures were harvested, and the cell pellets were frozen at −80°C until purification.

To lyse the cells, the pellets were slowly thawed on ice and resuspended in 4 mL of a Lysis Buffer containing 50 mM NaH2PO4, 300 mM NaCl, 10 mM imidazole, pH 8.0. Benzonase, Lysozyme (10mg/mL), and EDTA free protease inhibitor cocktail were further added to the Lysis buffer. The resuspension was kept on ice for two hours, followed by a heating step at 80°C for 15 min. The heated mixture was centrifuged at 8000 g, at 4°C for 20 min, and the supernatant containing the soluble fraction was collected and filtered sterilized before purification in the ÄKTA Pure chromatograph (GE Healthcare).

### Protein purification

PfuX7, Neq2X, and Neq2X7 were purified by affinity chromatography using a 1ml HisTrap™HP on an ÄKTA Pure chromatography system (GE Healthcare). After washing of the column, the target protein was eluted using a gradient protocol and the following buffers: wash buffer: 50 mM NaH_2_PO_4_, 300 mM NaCl, 20 mM imidazole (pH 8.0) and elution buffer: 50 mM NaH_2_PO_4_, 300 mM NaCl, 500 mM imidazole (pH 8.0). Peak fractions were analyzed on an InstantBlue (abcam) stained SDS-PAGE gel. Fractions containing the polymerase were then pooled, concentrated, desalted using a PD-10 desalting column (Merck), and loaded on a preparative Superdex200 increase 10/300 GL (GE Healthcare) column for size exclusion chromatography using a running buffer containing 20 mM Tris-HCl, 10 mM KCl, 6 mM (NH_4_)2SO_4_, 2 mM MgSO_4_ (pH 8.8). Peak fractions were analyzed on an InstantBlue (abcam) stained SDS-PAGE gel and subsequently pooled (see Supplementary Figure 2A). Samples were initially diluted 1:2 in storage buffer (20 mM Tris-HCl (pH 8.8), 10 mM KCl, 6 mM (NH_4_)2SO_4_, 2 mM MgSO_4_, 0.1 mg/mL BSA (nuclease free), 0.1% Triton X-100 and 50% glycerol) and subsequently stored at −20°C.

Protein concentration was determined by densitometry using the Fiji software^32^ as the BSA-containing buffer in the purified polymerase sample does not give accurate protein concentration measurement, and standard protein quantification methods could not be used. Triplicate representative sample preparations were loaded on an SDS-PAGE gel for each polymerase (PfuX7, Neq2X, and Neq2X7) containing 5 μL of the sample + 5μl of sample buffer. We used a serial dilution of a protein with a known protein concentration to estimate the protein concentration, which we used to calculate the standard curve, from which we figured the protein concentration of PfuX7, Neq2X, and Neq2X7 (see Supplementary Figure 2B, and C).

### DNA polymerase activity assay

To determine the polymerase activities of PfuX7, Neq2X, and Neq2X7, we followed the fluorescence method described by^33^ with minor modifications. In a 15 μL solution containing 1.2 pmol of the single-stranded M13mp18 DNA (Bionordika) previously annealed with 1.6 pmol of the primer UPlong (IDT) was mixed with 0.2 mM mixture of each dNTP (dATP, dCTP, dGTP, and dTTP), 2.5μL of 5xHF buffer (Thermofisher Scientific) and water to 15μL. We used three different DNA polymerase protein concentrations (0.5, 1, and 3 pmol) to determine the DNA polymerase activity, adjusted to a final volume of 5 μL. After ssDNA-primer-dNTPs-buffer temperature equilibration at 72 degrees, we added 5μL containing the polymerase, and the reaction was placed back at 72 degrees in a PCR cycler for 1, 3, and 15 minutes. Triplicate replicates were used of each concentration and time point for statistical analysis. The reaction was terminated by adding 1 μL of 0.5 mM EDTA, following the addition of 103.8 μL TE buffer (10 mM Tris-HCl, pH 7.9, 50 mM KCl, 1mM EDTA) and 0.2125 μL Pico488 dsDNA quantification reagent (Lumiprobe) to a final volume of 200 μL per sample. The 200 μL mixture was transferred to a 96-well black plate, and after 4-minute incubation, the absorbance was measured at 485 nm excitation and 528 nm emission with a top gain of 70. Double-stranded DNA of a known concentration measured by a Qubit 2.0 fluorometer (Invitrogen) were used to fit a linear regression to calculate the DNA amplification produced by the polymerases (Figure 1D, E, F)).

### PCR conditions

All PCR conditions unless stated otherwise were run using the following standard PCR protocol: DNA denaturation step of 98°C for 3 minutes, and 30 cycles of a second denaturation step at 98°C for 30 seconds, a primer annealing step of 60°C for 30 seconds, and an amplification step at 72°C with variable times. Primer sequences and templates are listed in Supplementary Table 1, 3, 4, and 5.

### Amplification of GC-rich DNA

Three templates Scat1, Scat2, and Tth2 of high GC content (see Supplementary Table 4), were chosen to test DNA polymerase performance. A touchdown PCR consisting of the following steps was used: an initial denaturation step of 98°C for four minutes, followed by 10 cycles of 98°C for 45 seconds, annealing at 65°C with one degree decrement every cycle, and an extension time of 1 minute, afterward followed by 30 cycles of 98°C for 45 seconds, annealing at 55°C, and extension of 1 minute. Primer pairs, templates and GC content is listed in Supplementary Table 4.

## Acknowledgments

We would like to thank Maja Lieven for her help input in DNA polymerase production and characterization, Timian Rindal for assisting in the laboratory, Folmer Fredslund for guiding us during the purification in the ÄKTA Pure chromatograph machine and finally, Viviënne Mol for providing us with the templates containing high GC DNA from *Streptomyces.*

