## Supplementary Materials for "Neq2X7: a multi-purpose and open-source fusion DNA polymerase for advanced DNA engineering and diagnostics PCR"

Addresses of all authors

Cristina Hernández Rollán:

Anja K. Ehrmann:

Morten H. H. Nørholm:

#### Supplementary Figure 1

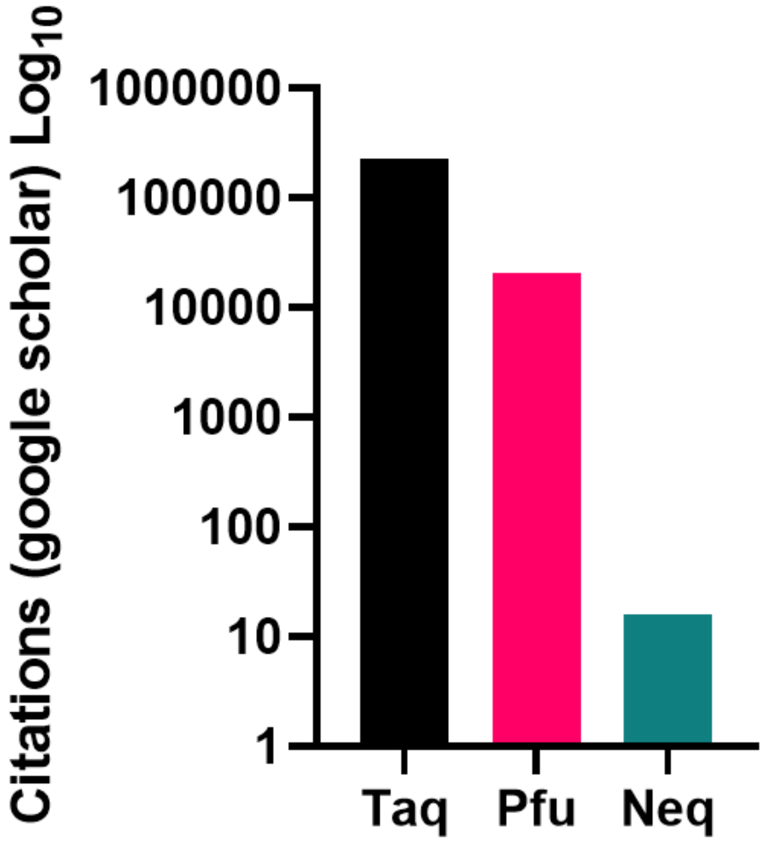


**Supplementary Figure 1.** Number of citations in Google scholar for the keywords Taq, Pfu, and Neq DNA polymerases represented on a log scale.

### Supplementary Figure 2

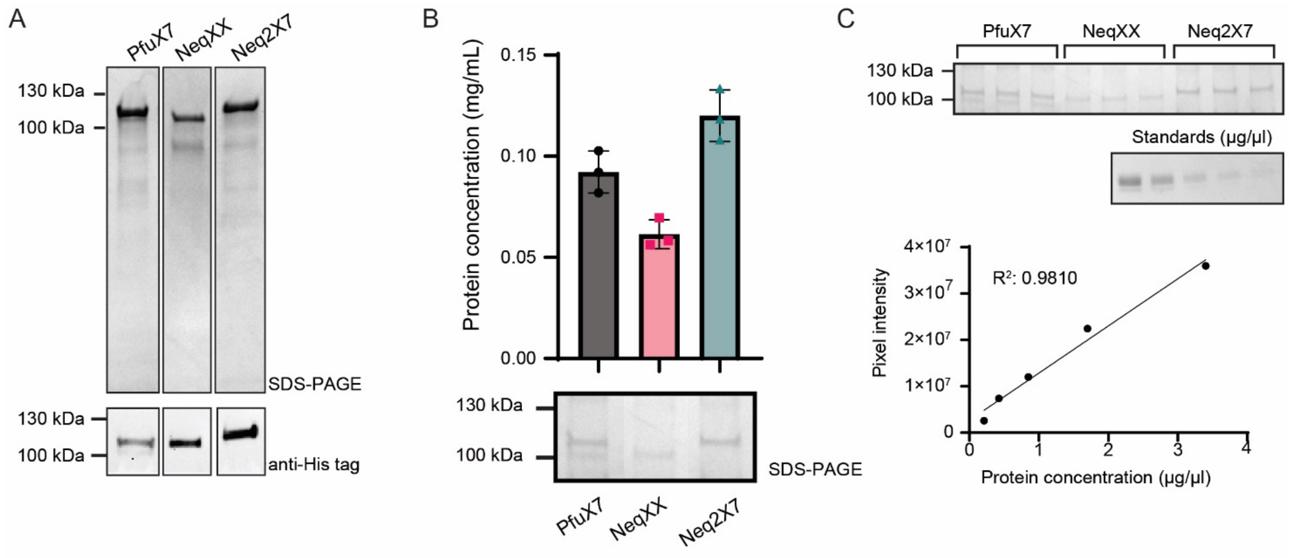


**Supplementary Figure 2. Estimation of the purity and concentration of the PfuX7, Neq2X, and Neq2X7 polymerases after Ni-NTA affinity and gel filtration purification**. (A) Coomassie blue-stained SDS–PAGE gel (upper panel) and Western blot (lower panel) of the three DNA polymerases after purification. (B) Protein concentration in mg/mL (upper panel) and a representative sample in an SDS-PAGE (lower panel). (C) SDS-PAGE (upper panels) with triplicate samples for each polymerase was used to calculate the protein concentration using the Fiji software[28]. The estimated concentrations were fitted to a regression line based on standards with known protein concentrations (lower panel). The molecular weights of the polymerases are 97.6 kDa for PfuX7, 95.7 kDa for Neq2X, and 103.2 kDa for Neq2X7.

### Supplementary Figure 3

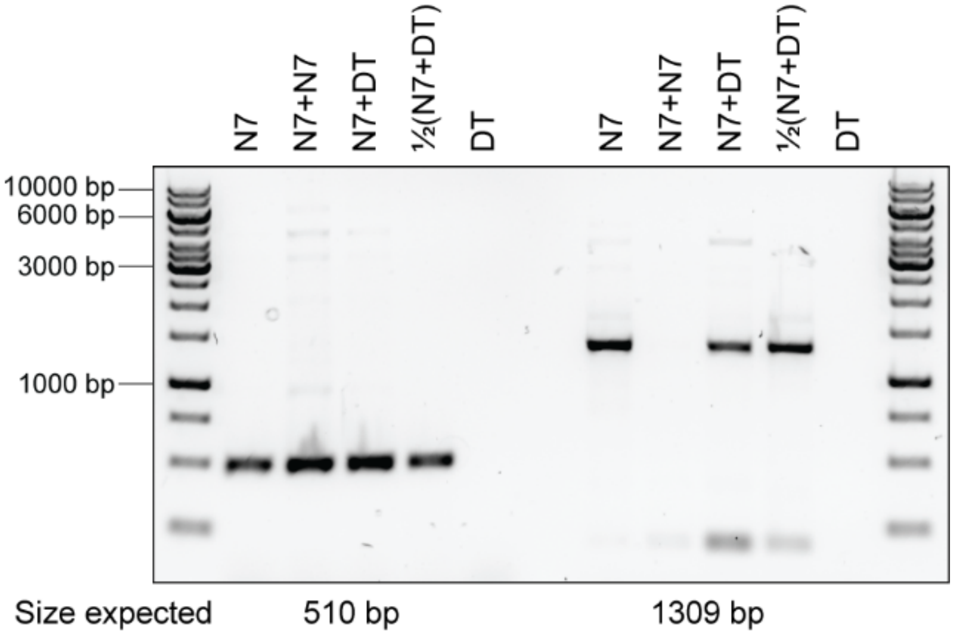


**Supplementary Figure 3. Evaluation of PCR performance of a mixture of Neq2X7 and DreamTaq DNA polymerases**. Two regions of the *E. coli* genome were amplified using either 1 pmol Neq2X7 (N7), 2 pmol Neq2X7 (N7 + N7), 1 pmol Neq2X7 + 0.625 U DreamTaq (Thermo Scientific) (N7 + DT), 0.5 pmol Neq2X7 + 0.31 U DreamTaq (½(N7+DT)) or 0.625 U DreamTaq (DT) in 25 µl reactions.

### Supplementary Table 1

| No. | Name | Sequence (5' --> 3') |
|---|---|---|
| 1 | 62 | TAATACGACTCACTATAGGGGAATTG |
| 2 | 63 | GCTCAGCGGTGGCAGCAGCCAACTCAGCTT |
| 3 | 4369 | AAATAAGAGAUCCGGCTGCTAACAAAGC |
| 4 | 4370 | ATCTCTTATTUAAAGAAATCTGTTAGTTTTTTGCCTGTGAATGTG |
| 5 | P341_Scat_U_FW_1 | AGGCGAUAGGAGGAATATACATGAGTCCTACGCCGCACA |
| 6 | P342_Scat_U_rev_1 | AGCCGUGCAGCACGGTCAGCGTGGT |
| 7 | 5868 | TTCCCAGTCACGACGTTGTAAAACGACGGCCAGTG |
| 8 | P343_Scat_U_FW_2 | ACGGCUCCAACCTGGGCGCCTGCCGGGA |
| 9 | P344_Scat_U_rev_2 | GGTGCGAUTCACCCGGCGGCGTACACGT |
| 10 | P346_Tth_U_FW_2 | AGAACCUCGAGGGCGACGAGCTGAAGA |
| 11 | P326_Tth_U_rev | GGTGCGAUTTAGTCAAAGACATCTGTGGCATACC |
| 12 | 4817 | AAACGCTGTCTTGGAACCTA |
| 13 | 4818 | AAACTGTCAGTTTTGGGCCAT |
| 14 | 4225 | CGTATGCATTGCAGACCTTGTGG |
| 15 | 4226 | GCACGATCCAACAGGCGAGC |
| 16 | 4229 | TATGGTGGTGAAGGGCGGTTC |
| 17 | 4230 | CGACGGTGATATTCCTCGCTC |
| 18 | 4703 | ACGGCTTCAGAATTTCTCAAGAC |
| 19 | 4926 | GCAAATGGCATTCTGACATCC |

**Supplementary Table 1. Oligonucleotides used in this study.**

### Supplementary Table 2

| Strain | Genotype | Source/Reference |
|---|---|---|
| *E. coli* NEB5α | *fhuA2* Δ(*argF-lacZ*)U169 *phoA glnV44* Φ80 Δ(*lacZ*)M15 *gyrA96 recA1 relA1 endA1 thi-1 hsdR17* | *a* |
| *E. coli* Rosetta BL21(DE3) | F-*ompT* *hsdS*$_B$($r_B$- $m_B$-) *gal dcm* (DE3) pLysSRARE2 ($Cam^R$) | *b* |

**Supplementary Table 2. Strains used in this study.**

[a]NEB, Ipswich, MA, USA.

[b]Novagen, Merck KGaA, Darmstadt, Germany.

### Supplementary Table 3

| Fragment | FW primer | RV primer |
|---|---|---|
| **3300 bp** | 4817 | 4929 |
| **6500 bp** | 4817 | 4703 |
| **12000 bp** | 4817 | 4818 |

**Supplementary Table 3. PCR-fragments.** Template was a pPIC9K derived plasmid.

### Supplementary Table 4

| Gene | FW primer | RV primer | GC content | PCR product |
|---|---|---|---|---|
| **Scat1** | 341 | 342 | 73% | 1540 bp |
| **Scat2** | 343 | 344 | 76.6% | 1680 bp |
| **Tth2** | 346 | 326 | 66.3% | 1516 bp |

**Supplementary Table 4. PCR products and primers used for GC amplification PCR.**

### Supplementary Table 5

| Step | Temperature | Time (seconds) | Cycle |
|---|---|---|---|
| **DNA denaturation** | 98°C | 40 | 1 |
| **DNA denaturation** | 98°C | 2 | 20 |
| **Annealing** | 60°C | 2 | |
| **Extension** | 72°C | 20 | |

**Supplementary Table 5. Fast PCR protocol used in this study.**
